# RNAiSpline: A Deep learning model for siRNA efficacy prediction

**DOI:** 10.64898/2026.02.14.705949

**Authors:** Sidhardha Reddy Surkanti, Vishnu Vardhan Kasturi, Sriram Shravan Saligram, Bhargava Chary Basangari, Vani Kondaparthi

## Abstract

RNA interference (RNAi) is a crucial biological post-transcriptional gene silencing mechanism where small interfering RNA (siRNA) guides RNA-induced silencing complex (RISC) to bind with messenger RNA (mRNA) thereby silencing it and stopping protein formation. We exploit this process to prevent the formation of harmful proteins by silencing mRNA before it is translated into protein through an effective siRNA. There exists a need to develop a computational model that predicts the effectiveness of siRNA on a given mRNA.

Designing a model is challenging, as the data availability is either scarce or biased, and existing models lack generalization ability, even though the parameters to training samples ratio is very high. To overcome these challenges, we introduce RNAiSpline, which incorporates self-supervised pretraining and fine-tuning with Kalmogorov-Arnold Network (KAN), Convolutional Neural Network (CNN), and Transformer Encoder.

Evaluation on the independent test dataset yields an ROC-AUC of 0.8175, an F1 score of 0.7717, and Pearson correlation of 0.6032, making RNAiSpline a robust model for siRNA efficacy prediction.

## 1. Introduction

RNA interference (RNAi) is a post-transcriptional gene silencing mechanism, a crucial part of both functional genomics research and therapeutic development [1]. In this process, small interfering RNA (siRNA), typically 21-23 nucleotides in length, is loaded into the RNA-Induced Silencing Complex (RISC), which contains Argonaute-2 (AGO2) protein. After the formation of this complex, siRNA duplex selects the strand based on thermodynamic asymmetry, where the strand with low base pairing stability at its 5’ end is loaded as the guide strand into RISC [2]. This RISC-siRNA complex recognises the complementary messenger RNA (mRNA) sequence through Watson-Crick base pairing, leading to endonucleolytic cleavage and degradation of the target mRNA, thereby preventing protein translation. While RNAi is a powerful tool for studying gene functions and developing therapeutic applications, the design of highly efficient siRNA remains a critical bottleneck, as efficacy varies drastically depending on sequence-specific features, thermodynamic properties, and structural accessibility of both siRNA and target mRNA

The first computational approaches that appeared in 2004, were based on empirical observations from small experimental datasets in order to derive sequence-specific rules. Reynolds examined 180 siRNAs and identified eight features in their sequences. Ui-Tei [3] had developed four strict selection criteria that focused on A/U nucleotides. Amarzguioui [4] had incorporated thermodynamic asymmetry and identified functionally essential motifs. However, these rule-based algorithms were inherently limited in modeling complex non-linear interactions and synergistic effects among multiple sequence determinants.

With the development of machine learning techniques to extract complex patterns from large-scale experimental datasets, the computational landscape changed further. Huesken initiated this change in 2005 with BIOPREDsi [5], which utilizes artificial neural networks trained on 2431 siRNAs having quantitative efficacy measurements. Ichihara proposed i-Score [6] in 2007 as a simplified linear regression approach using only position-specific nucleotide preferences. Lu and Mathews proposed OligoWalk [7] in 2008, which introduced a thermodynamics-based approach that applies partition function calculations for predicting free energy changes occurring during siRNA-mRNA hybridization. Vert extended the linear modeling approach in 2012, proposing DSIR [8], which combined position-specific nucleotide preferences with particular motifs on the guide strand. Qureshi filled an important void in 2013 with VIRsiRNApred [9], having proposed the first virus-specific efficacy prediction algorithm by applying support vector machines trained on 1725 experimentally verified viral siRNAs targeting 37 human viruses. Dar advanced the field further in 2016 with SMEpred [10], the first predictor for chemically modified siRNAs addressing 30 commonly used chemical modifications, trained on 3031 chemically modified siRNA sequences. However, these machine learning algorithms still relied on handcrafted feature engineering and had difficulties in learning hierarchical representations from raw sequence information directly.

With the emergence of deep learning, deep neural architectures started to represent features hierarchically and without explicit feature engineering. The development of convolutional neural networks [11] around 2017 and 2018 explored capturing local sequence motifs and position patterns through learnable convolutional filters. In 2022, for the first time in this domain, La Rosa introduced graph neural network methodology with GNN4siRNA [12], which fundamentally reconceptualized efficacy prediction as modeling siRNA-mRNA interactions within graph-structured representations. GNN4siRNA utilized a modified GraphSAGE framework [13] for encoding siRNAs as 3-mer features and mRNAs as 4-mer features in order to process node and edge features. A 3-mer (or trigram) represents all possible sequences of 3 consecutive nucleotides from the RNA sequence. Similarly, a 4-mer represents all possible sequences of four successive nucleotides. However, it performed very poorly when both the training and test datasets were drawn from different distributions of data points, limiting cross-dataset applicability. In 2024, Long developed another architecture, siRNADiscovery [14], which is a comprehensive model embedding empirical features from established design principles and non-empirical features learned from data patterns. Despite such remarkable achievements, GNNs [15] introduce considerable computational complexity, which raises serious overfitting concerns using small-scale training data and may pose practical deployment barriers.

Recent models have adopted a transformer architecture [16], leveraging its established performance in modeling biological sequences. Bai et al. developed OligoFormer [17], with an elaborate multi-module architecture consisting of thermodynamic feature calculation, RNA-FM [18], a pre-trained transformer encoder-only foundation model for RNA sequence, and OligoEncoder that incorporated CNN layers to capture local motifs, bidirectional LSTM [19] for sequential dependencies, and transformer encoder layers for long-range interactions. Comprehensive integration of multiple information sources enabled OligoFormer to achieve unparalleled performance on benchmark tests while demonstrating solid generalization to a wide range of experimental conditions.

Despite years of methodological evolution, several important challenges remain that critically hinder the prediction of siRNA efficacy. One of the main factors in predicting siRNA efficacy is dataset bias, limited training data and heterogeneity of data. Second, a proper aggregation for the combination of extracted features from the sequence.

To address these challenges, we introduce RNAiSpline, a novel deep learning framework including KANs [20], convolutional neural networks, and a transformer for siRNA efficacy prediction. Instead of standard Multi-Layer Perceptrons with fixed node activation functions, KAN uses learnable univariate functions on edges based on the Kolmogorov-Arnold representation theorem, which that states any multivariate continuous function can be represented as a superposition of continuous one-dimensional functions and addition operations.

RNAiSpline addresses the critical limitations of existing approaches. First, with only 955,968 parameters, our model achieves competitive performance without relying on pre-trained models for embeddings. This architecture effectively integrates CNN layers for local motif extraction, transformer encoders for capturing long-range sequence dependencies, and KAN-based classification layers with Cox-de Boor B-splines [21] for final efficacy prediction. Second, processing one-hot-encoded sequences together with thermodynamic features directly enables RNAiSpline to learn complex, nonlinear decision boundaries. B-spline activations within KAN layers yield smooth, visualizable functions that interpret how specific sequence patterns contribute to efficacy predictions. Third, the framework includes self-supervised pre-training on sequence reconstruction tasks, where the model learns generalizable representations directly from unlabelled siRNA-mRNA pairs before fine-tuning for efficacy prediction. Finally, this lightweight architecture enables fast inference on standard hardware, making RNAiSpline practical for high-throughput siRNA efficacy prediction.

RNAiSpline enables a wide range of applications in RNAi research and therapeutic development by bringing together the representations learned by state-of-the-art deep learning architectures with the efficiency and interpretability of Kolmogorov-Arnold Networks implemented through B-spline bases. This work shows that, through a responsible architectural design driven by biological principles, competitive performance can be achieved without resorting to massive pre-trained models.

## 2. Methodology

### 2.1. Datasets and preprocessing

#### 2.1.1. Datasets

A diverse array of datasets is collected for the accuracy and robustness of our model. A total of nine datasets of 3,714 siRNAs and 75 mRNAs from previous studies, including Huesken [5], Takayuki [22], Amarzguioui [23], Haborth [24], Hsieh [25], Khvorova [26], Reynolds [27], Vickers [28], and Ui-Tei [3] are used. Datasets are divided into three sets: the Huesken dataset, the Takayuki dataset, and Mixset by combining all remaining datasets. The efficacy of siRNAs in all datasets is normalized, ranging from 0 to 100%, and 70% of the maximum efficacy was used as the threshold to classify effective and ineffective siRNAs. 2361 samples from Huesken are used for training, 20% of 2361 is used for validation, Takayuki and Mixset are used for evaluating the model.

Table 1 presents Huesken, Takayuki and Mixset datasets used as siRNA sources.

**Table 1.** Various sources for siRNA datasets.

| Source | siRNA | mRNA | Cell Line | Dataset |
| --- | --- | --- | --- | --- |
| Huesken | 2431 | 34 | H1299 | Huesken |
| Takayuki | 702 | 1 | HeLa | Takayuki |
| Amarguioui | 46 | 4 | HaCaT | Mixset |
| Harborth | 44 | 1 | HeLa |  |
| Hsieh | 108 | 22 | HEK293T |  |
| Khvorova | 14 | 1 | HEK293 |  |
| Reynolds | 240 | 7 | HEK293 |  |
| Vickers | 76 | 2 | T24 |  |
| Ui-Tei | 53 | 3 | HeLa |  |

##### The Huesken Training Dataset

The Huesken training dataset contains 2,361 siRNAs generated by Huesken et al. using standardized high-throughput dual-luciferase reporter assays. All experiments were conducted on H1299 human non-small cell lung carcinoma cells at a consistent 100 nM siRNA concentration. This dataset shows a nearly normal efficacy distribution with a mean knockdown efficiency of 0.51(standard deviation), minimal skewness of −0.064, and mean GC content of 50.37%.

##### The Mixset Testing Dataset

In contrast, the Mixset testing dataset aggregates 472 siRNAs from multiple independent studies. The experiments used diverse cell lines (HeLa, HaCaT, HEK293, HEK293T, and T24) and employed various methodological approaches, including quantitative RT-PCR [29], Western blot protein quantification [30], and different reporter gene assays with varying normalization protocols. The siRNA concentrations also differed substantially across studies: Reynolds used 100 nM, Harborth used 10 nM, and Hsieh and Vickers used 20-50 nM. This heterogeneous dataset exhibits a different efficacy distribution with a mean of 0.69 and a negative skewness of −0.770, indicating enrichment toward highly effective siRNAs. The mean GC content is 49.48%.

##### Statistical Differences between Datasets

Statistical analyses reveal significant divergence between the two datasets across multiple dimensions. Independent t-tests show dramatically different efficacy means (p=5.67×10−78), while Levene’s test [31] confirms unequal variances (p=4.02×10−69). The Kolmogorov-Smirnov [32] test demonstrates different distribution shapes (p=2.98×10−90), and GC content distributions also differ significantly (p=4.55×10−2).

The concentration differences warrant particular attention. Higher concentrations like 100 nM can mask sequence-specific effectiveness through saturation effects and off-target modulation. Lower concentrations between 3-30 nM provide better discriminatory power for distinguishing highly functional siRNAs from moderately functional ones, since efficacy measurements follow dose-response relationships.

By training the model on the homogeneous Huesken dataset and testing it on the heterogeneous Mixset, we can evaluate how well the model generalizes across real-world variations. These include cell-line-specific biological differences, diverse experimental methodologies, and concentration-dependent effects that researchers encounter when working with siRNA efficacy data.

#### 2.1.2. Sequence processing

The datasets consist of variable siRNA and mRNA length. Many siRNA samples are of length 19 nucleotide and mRNAs of different length, to make it consistent and easy for model to extract features, siRNA of length 21 nucleotides was processed into 19 nucleotides by removing 2 overhangs at the 3’ end of the sequence and mRNA into 19 nucleotides to match siRNA length. The binding site of mRNA which is 19 nucleotides long is extracted with the reverse complement of siRNA.

#### 2.1.3. Thermodynamic features

RNAiSpline extracts 24 thermodynamic features that describe the siRNA interactions with its target mRNA and loads into the RISC complex. Thermodynamic features include Gibbs free energy and enthalpy values for all possible dinucleotide pairs, terminal AU pairs, and symmetry, the end differential free energy (D).

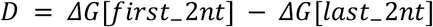

Where, first_2nt is first two nucleotides and last_2nt is the last two nucleotides of the sequence, gives a measure over the 5’ vs 3’ end energy, free energy at positions that have important functional roles, position 2-3 for seed region stability, position 12-13 for central duplex integrity, and position 17-18 for 3’ end stabilization, U/G/A/C content for binding strength, local stability patterns by calculating the occurrence frequency of GG, UA, CC, GC, UU between positions 0 and 17. Binary indicators are used to mark the occurrence of nucleotides at specific positions that are important for silencing, this includes position 1 for RISC loading, position2 for seed region recognition, and position 19 for overhang stability.

Thermodynamic parameters represent the fundamental physical properties of RNA that help us to understand how siRNA functions. The lower the ΔG values, the more stable are the interactions, meaning that binding is stronger. The difference in free energy within the two ends of siRNA is particularly important, when the 5’ end is less stable, the antisense strand is more likely to get loaded into the RISC complex which is crucial for effective gene silencing. RNAiSpline combines these 24 thermodynamic features with sequence patterns learned through convolutional and transformer layers. This allows the model to learn how the siRNA duplex forms and whether the target site on the mRNA is accessible. By analyzing sequence composition, structural stability, and their relationship to silence effectively, the model can make more accurate predictions than methods only considering the sequence itself.

### 2.2. The architecture of RNAiSpline

RNAiSpline comprises two stages of training: self-supervised pretraining and supervised fine-tuning for efficacy prediction. Pretraining compensates for the lack of labeled siRNA efficacy datasets. Thus, pretraining allows the model to learn sequence characteristics by only using unlabeled data. The self-supervised pretraining consists of a sequence reconstruction task, for each sample, the siRNA and mRNA sequence have their nucleotides randomly masked (with a masking probability of 15%) across their lengths. Three different masking strategies are performed on these masked locations: 80% of the locations are completely masked out and substituted by zero vectors, 10% of the locations are substituted with random nucleotides from (A, U, G, C) for noise and 10% remain the same. The masked siRNA and mRNA sequence are pre-trained separately using a CNN-Transformer architecture as shown in Figure 1.

**Fig. 1.**
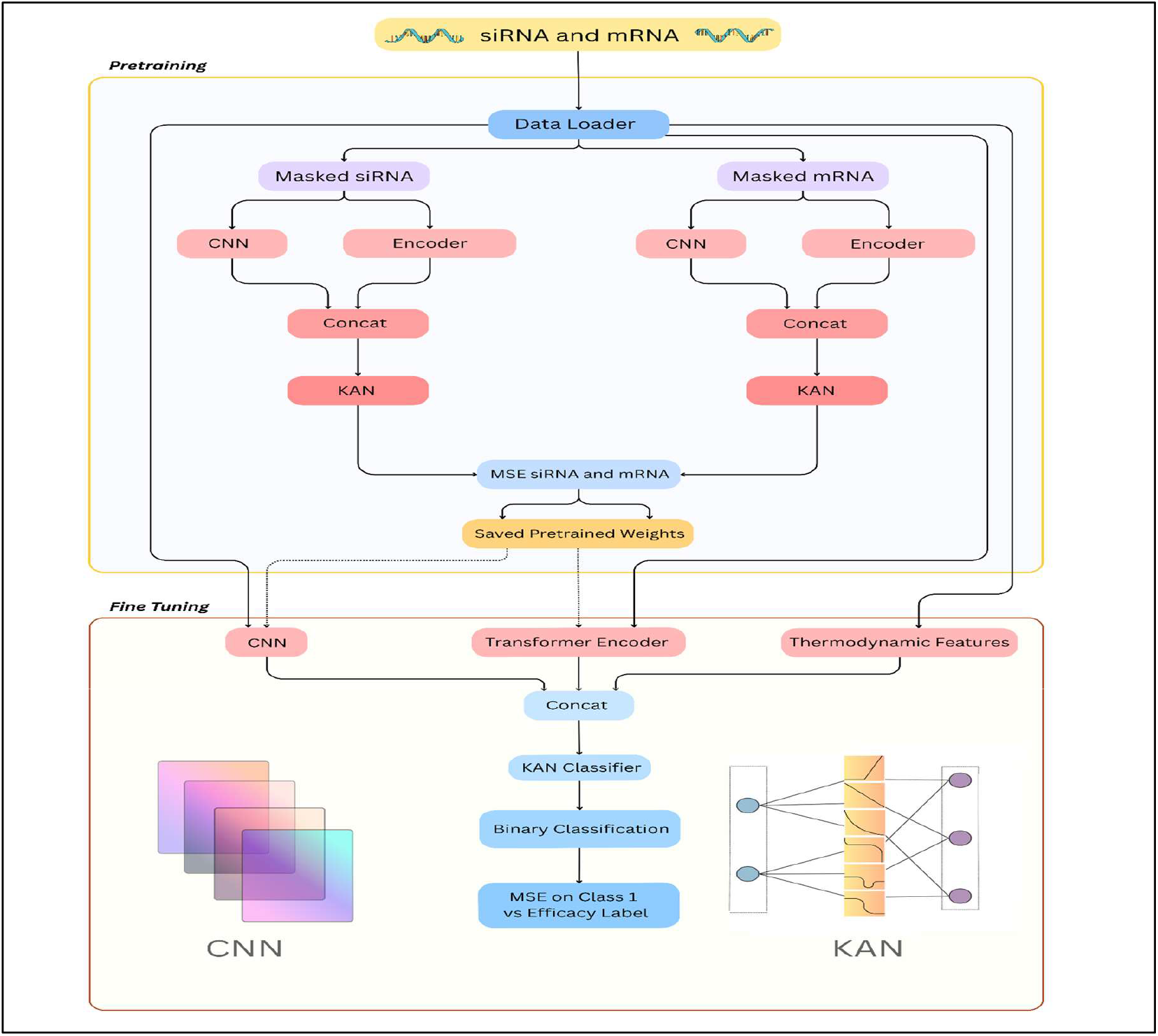
RNAiSpline architecture. Figure 1 projects the pretraining phase loads sequence, performs sequence regeneration task and saves base model weights of 1D CNN and Transformer. Fine-tuning phase loads sequence and base model weights to perform classification task.

CNN captures local sequence motifs, the Transformer captures positional dependencies and long-range interactions. The 96-dimensional features from the CNN branch and 64-dimensional features from the Transformer branch are combined into a 160-dimensional representation for both siRNA and mRNA. Each component of this fusion is input into a separate KAN reconstruction module with a specified architecture, where each reconstructor learns features and projects them back to the original sequence (19 × 4 bases). Pre-training uses mean squared error loss between reconstructed and original one-hot encoded sequence at masked positions only:

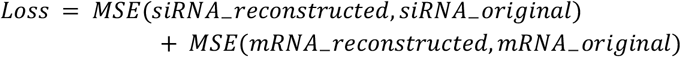

Since the reconstruction is trained together, the model learns unique features of each sequence and implicit siRNA-mRNA interactions. Pretraining provides robust weight initialization for the fine-tuning phase, enabling faster convergence and reducing the risk of overfitting when trained on small labelled datasets.

After pretraining, the learned weights from siRNA CNN layers, mRNA CNN layers, siRNA Transformer, and mRNA Transformer are saved and loaded into the final classification model. The KAN-based reconstruction modules used during pretraining are discarded and not transferred to the fine-tuning stage, as their purpose is only to reconstruct sequences. The fine-tuning architecture retains the identical dual-pathway CNN and Transformer modules described in the pretraining phase, but replaces the reconstruction-focused KAN modules with a classification-focused KAN to predict efficacy. The CNN branch contains a three-layer stacked convolutional architecture with 96 output channels. Every layer performs a 1D convolution with a kernel size of three and a padding of one to maintain sequence length, followed by ReLU [33] activation that introduces non-linearity. With this stacked CNN, the model can progressively learn patterns: the earlier layers capture simple motifs such as dinucleotide patterns while deeper layers identify complex multi-nucleotide patterns. After the convolutional layers, adaptive max pooling collapses the dimension to a 96-dimensional feature vector. The transformer branch captures long-range dependencies across the sequence, and the encoder uses eight attention heads and 128-dimensional feedforward layers. Multi-head self-attention calculates the scaled dot-product attention along with query, key, and value projections to learn positions that are most relevant to each other. Finally, adaptive average pooling reduces it into a single 64-dimensional vector that has the global contextual relationships uncovered by the transformer.

The combined 344-dimensional feature vector contains 24 dimensions of thermodynamic stability matrices, 96 from siRNA CNNs, 64 from siRNA transformer encoders, 96 from mRNA CNNs, and 64 from mRNA transformer encoders, and is an input to KAN classifier. Each KAN layer uses cubic B-splines with spline orders 3 and 5 grid points spanning -2 to 2, with the Cox-de Boor recursive algorithm efficiently computing basis functions while handling knot boundaries and edge cases. Spline weights from a three-dimensional tensor (out_dim × in_dim × num_basis), allowing each input-output connection to learn its unique spline transformation. The layer output combines these spline-based transformations with linear projections (spline_output × linear_output), enabling the network to capture both smooth non-linear patterns and direct linear dependencies simultaneously. The sequence *t*_*i*_ represents the positions where the polynomial pieces join (knots) and *k* determines the polynomial degree (linear, quadratic etc.,). The input *x* is the value at which the basis is evaluated.

1. Base Case: Piecewise Constant (Degree 0)

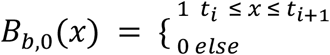
2. Recursive Case: Higher Degree *k*

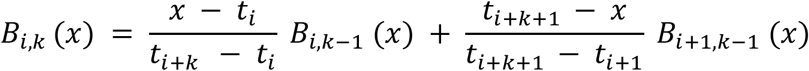

The network contains 774,912 trainable parameters, with 83.3% allocated to spline coefficients and 16.7% to linear weights, allowing for flexible function approximation while maintaining computational efficiency. The learned activation functions of layer 0, layer 1, layer 2, and layer 3 are represented as Figure 2, Figure 3, Figure 4 and Figure 5 respectively.

**Fig. 2.**
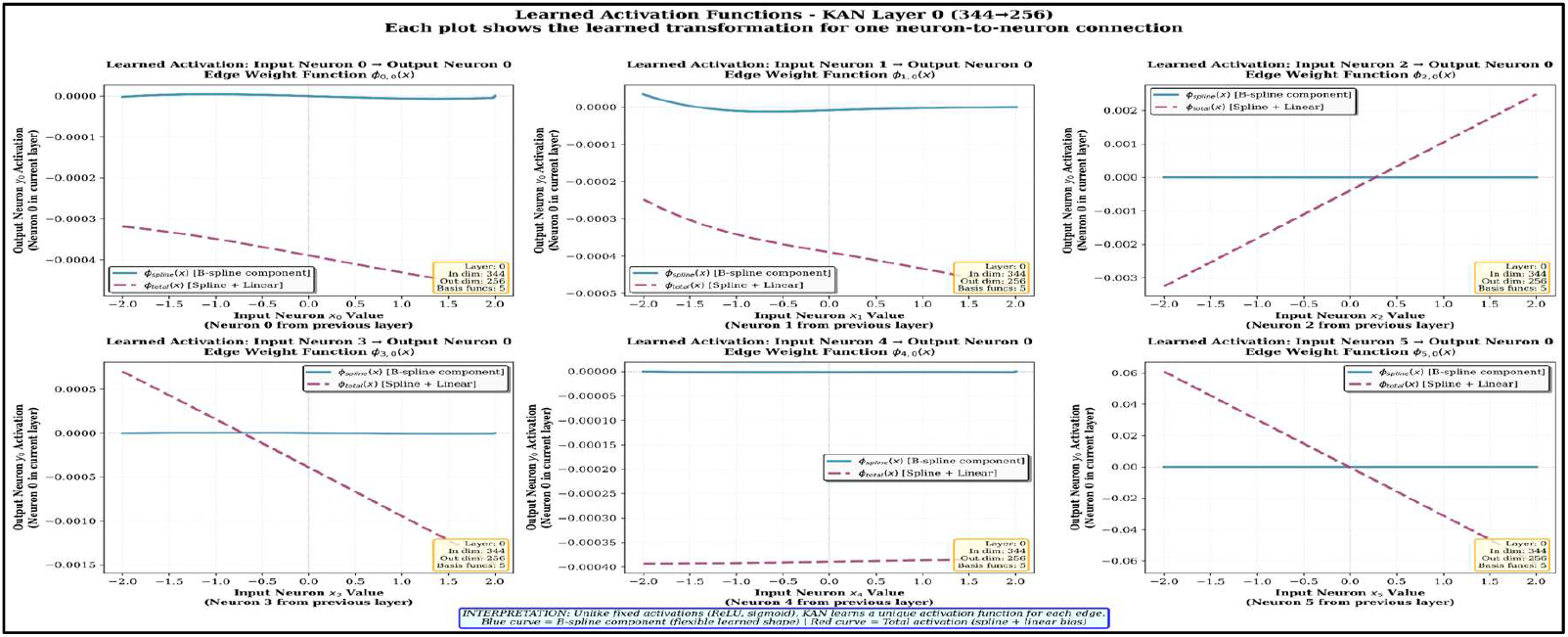
Layer 0: Input Feature Processing. Kan Layer 0 (344 to 256) learned activation functions is presented as line graph in Figure 2.

**Fig. 3.**
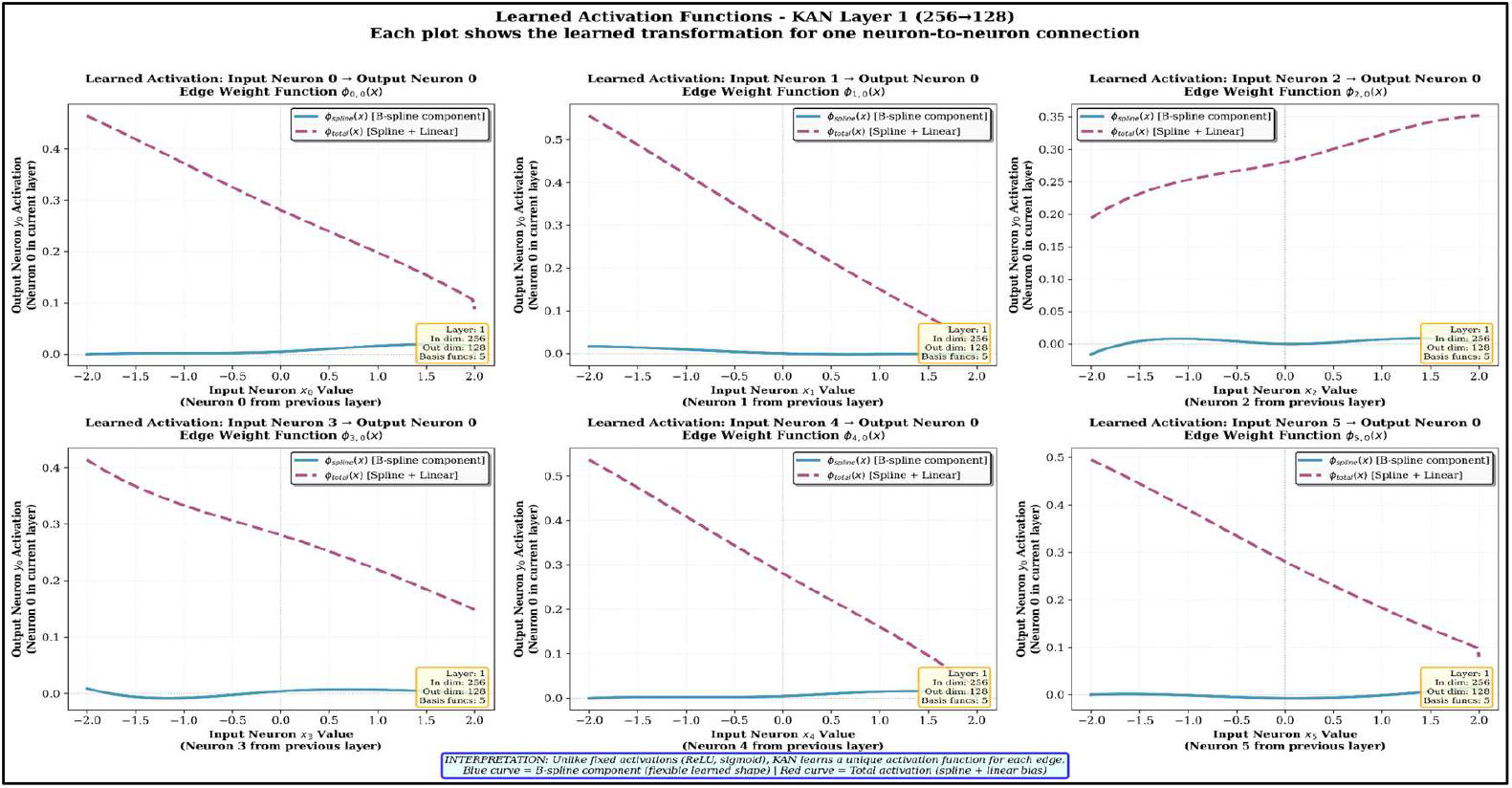
Layer 1: Intermediate Representation Refinement. KAN Layer 1 (256 to 128) learned activation functions are represented as line graph in Figure 3.

**Fig. 4.**
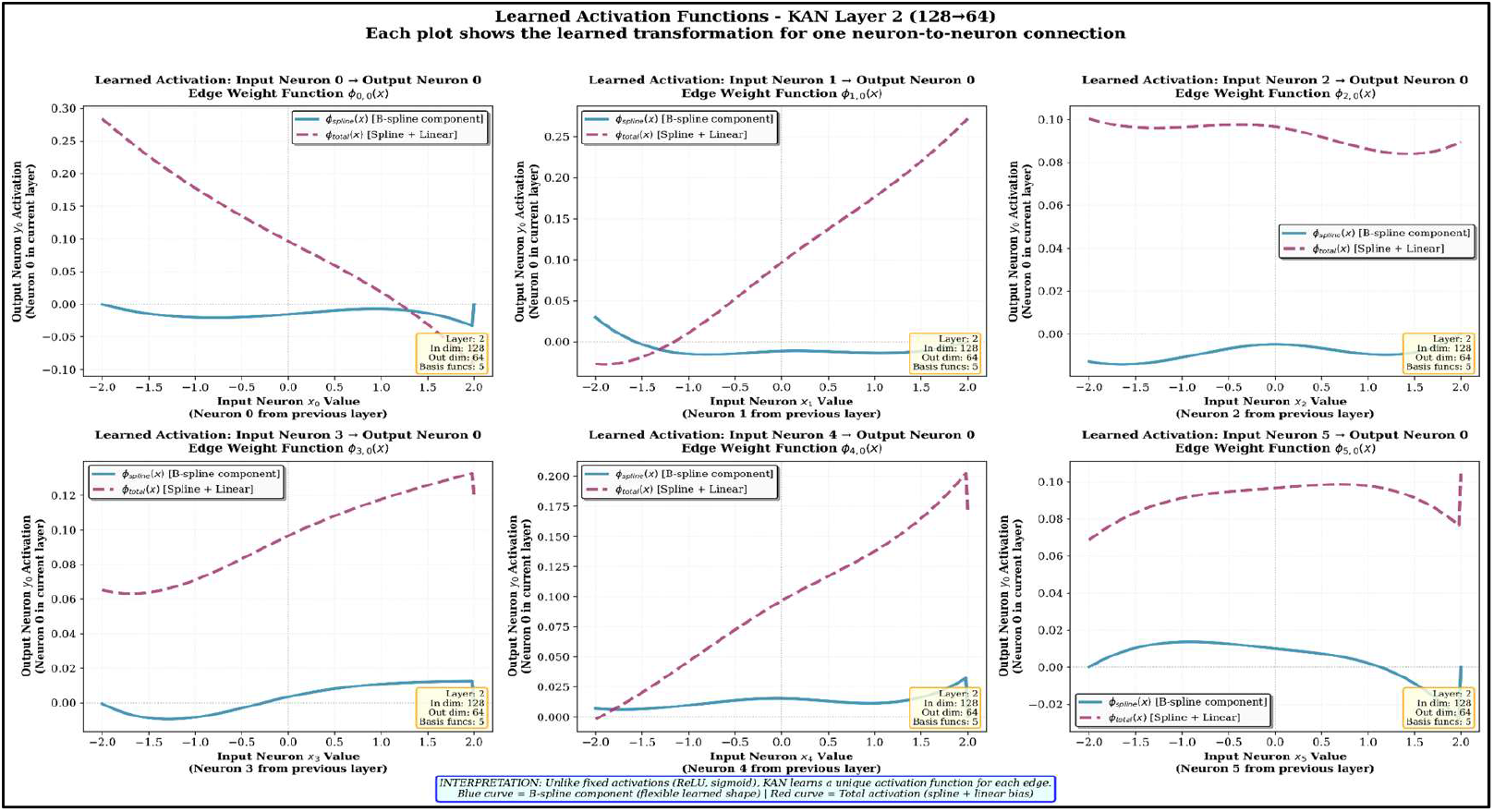
Layer 2: Deep Feature Abstraction. KAN Layer 2 (128 to 64) learned activation functions are depicted as graph in the Figure 4.

**Fig. 5.**
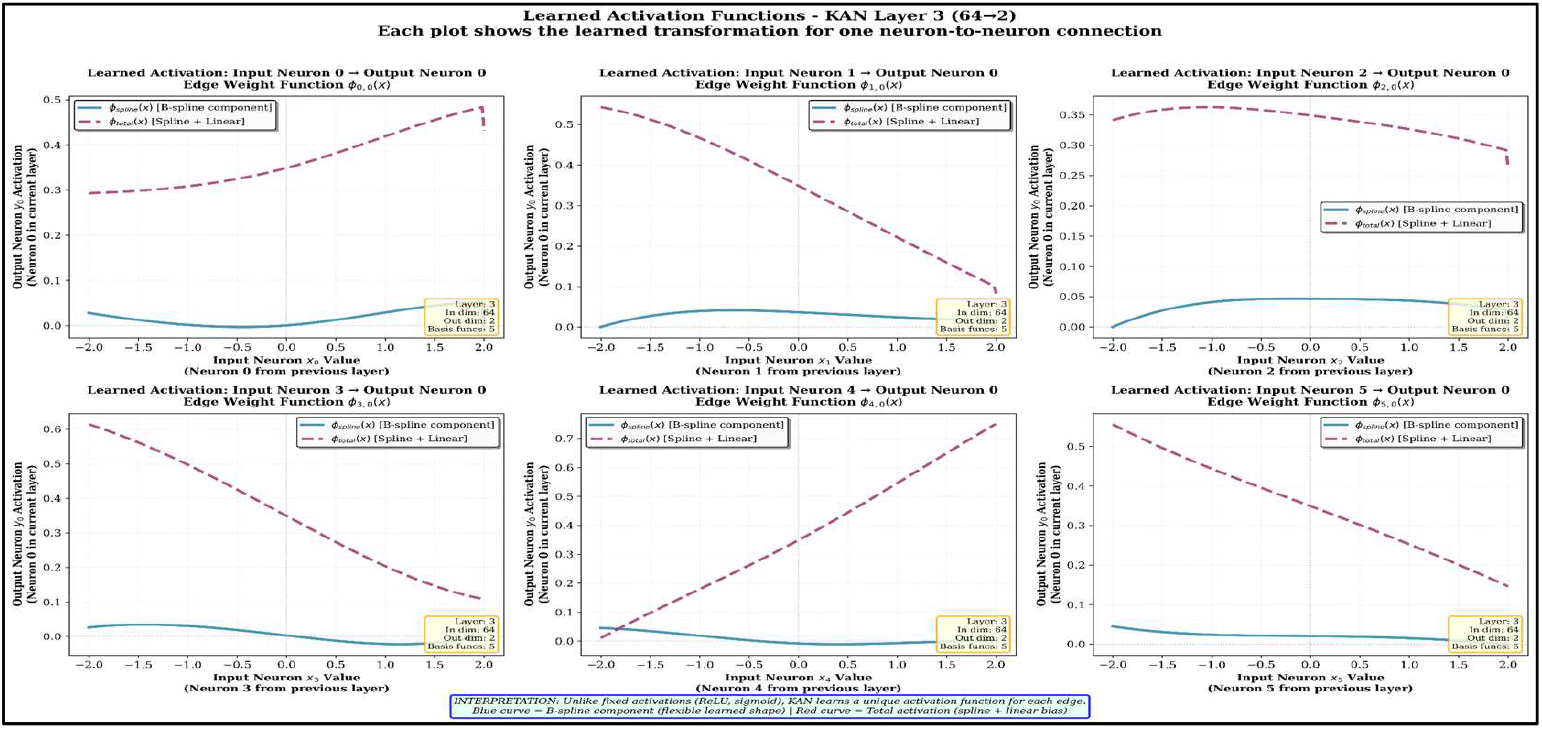
Layer 3: Classification Output. KAN Layer 3 (64 to 2) learned activation functions are shown in Figure 5.

The first KAN layer (layer 0) transforms 344 input features into 256 intermediate representations through 88,064 connections, each parameterized by six values (five spline coefficients and one linear weights). This layer contains 528,384 parameters, making it the most computationally intension component of the network. The learned spline weights exhibit a narrow distribution (mean: 0.000172, standard deviation: 0.005840), indicating selective activation patterns, while linear weights show broader variation (standard deviation: 0.028146), suggesting diverse feature combinations. The B-spline configuration with a knot vector [-2,-2,-2,-2,0,2,2,2,2] ensures *C*^l^ continuity, enabling smooth gradient flow during backpropagation.

The second layer (layer 1) reduced dimensionality from 256 to 128 neurons using 196,608 parameters across 32,768 connections. The spline weights (standard deviation: 0.010503) demonstrates increased variability compared to Layer 0, reflecting more specialized feature extraction. Linear weights show substantially higher variation (mean: 0.000944, standard deviation: 0.087262), with a median of 0.000147 indicating asymmetric weight distribution. This layer serves as a critical bottleneck, compressing high-dimensional representations while preserving discrimination information through learned non-linear transformations.

Third Layer (layer 2) further compresses representations from 128 to 64 neurons with 49,152 total parameters. The spline weights exhibit the highest variability observed (standard deviation: 0.018189), suggesting complex nonlinear feature interactions. Linear weight statistics (mean: 0.000643, standard deviation: 0.096916, median: 0.000704) reveal near-zero-centered distributions with significant spread, indicating balanced positive and negative contributions. The reduced parameter count (8,192 connections) enforces aggressive feature selection, extracting only the most salient patterns for final classifications.

The final layer (layer 3) projects 64 abstract features onto two output neurons through 768 parameters (640 spline, 128 linear). Spline weights maintain tight distributions (standard: 0.029639) despite reduced dimensionality, whereas linear weights show notable bias (mean: 0.005009, median: 0.014809), indicating systematic positive weighting toward specific class predictions.

B-spline basis functions offer distinct advantages over standard activation functions like ReLU or sigmoid. They provide smooth differentiability for stable gradient flow during backpropagation, local support ensuring each basis function affects only a limited input range without causing unintended global disturbance, and interpretability through visualization of learned spline shapes that reveal how specific input features influence predictions. This KAN-based design naturally aligns with the continuous nature of biological sequence relationships, in which minor sequence variations typically produce gradual efficacy changes rather than abrupt transitions.

## 3. Results and discussion

### 3.1. Evaluation

#### 3.1.1. Intra-dataset evaluation

RNAiSpline was trained on the Huesken, Mixset, and Takayuki datasets to assess its performance consistency across different experimental conditions and evaluated on each dataset separately. On the Huesken, RNAiSpline achieved 0.8538 ROC-AUC, and an F1 score of 0.7867 with a PCC of 0.6740. On the Mix dataset that combines multiple heterogeneous datasets including Amarzguioui, Harborth, Hsieh, Khvorova, Reynolds, Vickers, and Ui-Tei, RNAiSpline achieved the strongest performance in this paper with ROC-AUC of 0.8641, F1 score of 0.7791, and PCC of 0.7368. On the Takayuki dataset, RNAiSpline showed exceptional generalization with ROC-AUC of 0.8791, an F1 score of 0.5601, and a PCC of 0.7904; it significantly outperformed all baseline methods, including OligoFormer. These results indicate that the KAN-based architecture of RNAiSpline with Cox-de Boor B-splines effectively models complex sequence-efficacy relationships and is particularly good at correlation-based metrics reflecting quantitative prediction accuracy (Supplementary Table S1).

#### 3.1.2. Inter-dataset evaluation

To evaluate cross-dataset generalization capability, RNAiSpline was trained only on the Huesken dataset and tested on the Mix dataset. RNAiSpline achieved ROC-AUC of 0.8175, F1 score of 0.777 and Pearson correlation of 0.6032, surpassing baseline models including Monopoli-RF [34], OligoWalk [7], siRNAPred [35], i-Score [6], s-Biopredsi, DSIR [36] and OligoFormer [17]. Metrics of all models were taken from the OligoFormer research paper.

The metrics available in Table 2 as per OligoFormer research paper and tested on inter dataset after training it on Huesken dataset. The metrics highlighted in bold is the best obtained value among the models for specified parameters.

**Table 2.**
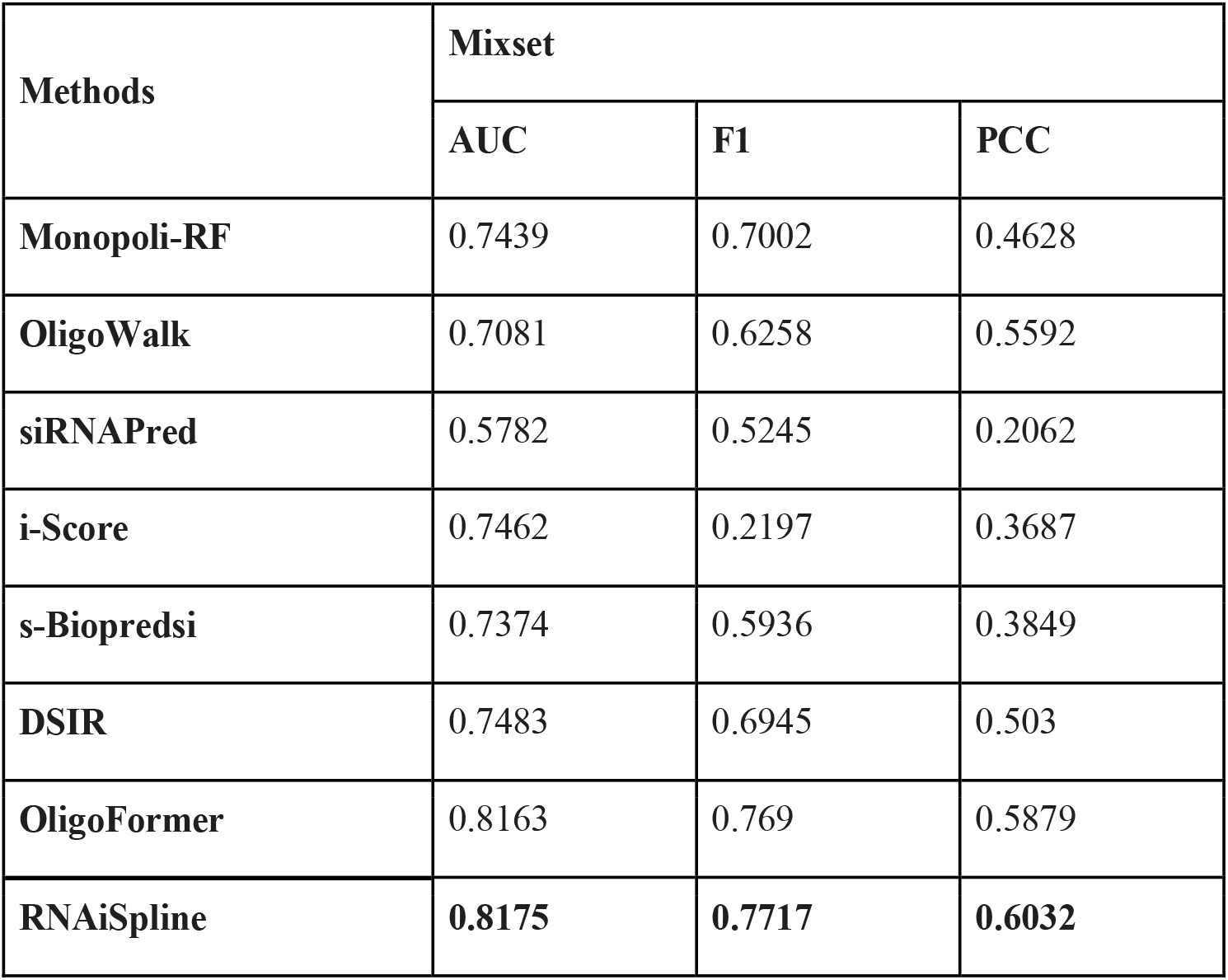
Inter-dataset metric for testing.

Inter dataset evaluation reveals that RNAiSpline performs better compared to existing methods, with AUC, F1-score, and PCC of 0.8175, 0.7717 and 0.6032 respectively as given in Table 2. The improved PCC metric demonstrates that the model’s predictions exhibit stronger correlation with experimental efficacy measurements compared to baseline approaches.

### 3.2. Ablation study

To evaluate the performance of each component in RNAiSpline, Ablation study was conducted by systematically removing or replacing individual components while keeping all other components as it is. Each ablation involved training the architecture with identical hyper parameters, data splits and methods to ensure the same comparison. Five ablations were done, replacing the KAN classifier with a standard Multi-Layer Perceptron, removing the transformer encoder layers, excluding thermodynamic features, removing CNN layers and training without the self-supervised pretraining phase.

Ablation experiments depicted in Figure 6 as a line graph demonstrate that replacing the MLP classifier with KAN improves AUC-ROC performance, while pretraining proves critical for PCC optimization. Thermodynamic features exhibit the most substantial impact on the F1 score.

**Fig. 6.**
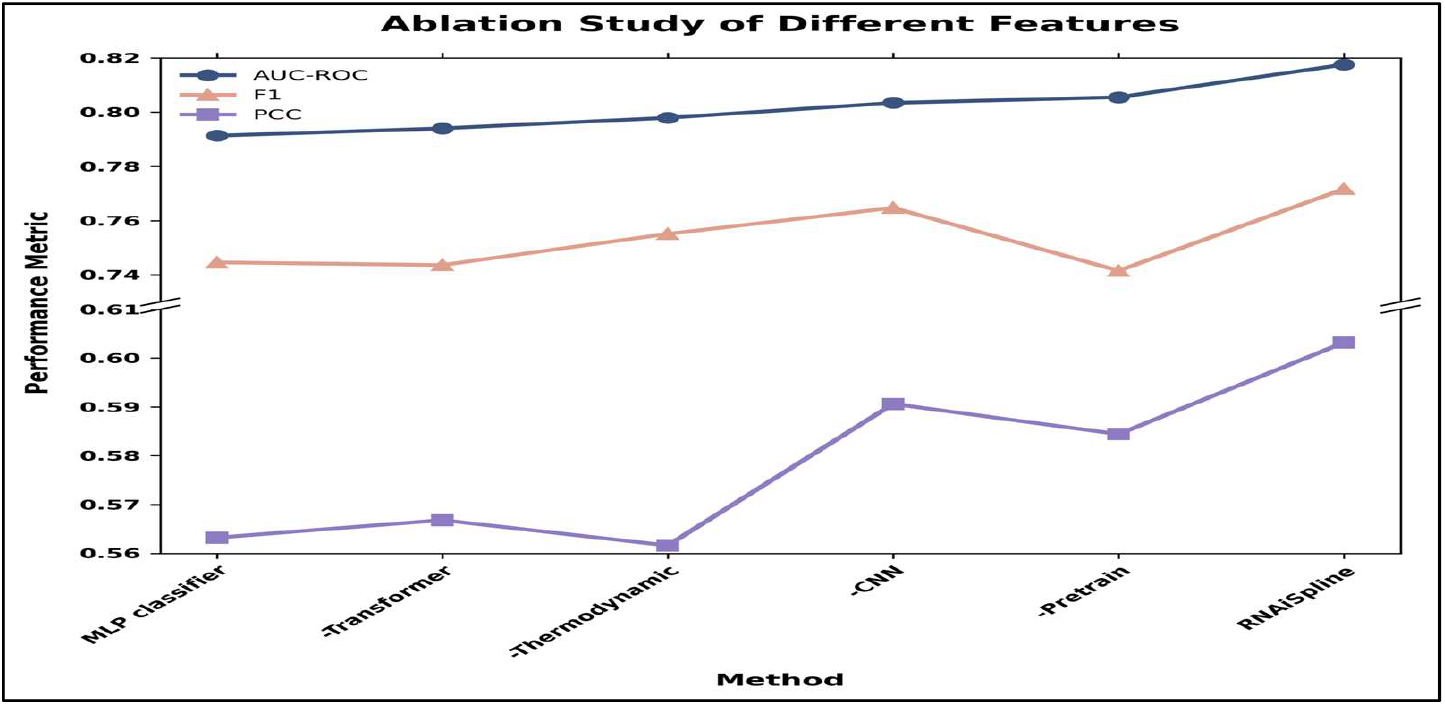
Ablation study of different features of RNAiSpline. The MLP classifier is used instead of the KAN classifier, and the rest is removed while keeping everything constant.

## 4. Conclusion

RNAiSpline demonstrates that a well-designed architecture can achieve competitive results without relying on external pretrained embedding models or transfer learning. By integrating supervised learning, the model can partially overcome dataset limitations while simultaneously acquiring the ability to understand sequences. By integrating CNN for local motifs, Transformer for long-range dependency, and Cox-de Boor B-Spline KAN classifiers with thermodynamic features, the model achieves superior performance on unseen data with 0.8175 AUC, 0.7717 F1-Score, and 0.6032 PCC.

However, there is still room for improvement. There is a severe deficit of large, high-quality public datasets. If there are more public datasets in the future, the model can achieve better metrics while maintaining or increasing its generalization ability. The Cox-de Boor B-Spline basis function calculation is a sequential recursive function. To further improve the computational speed, MatrixKAN uses matrix representation of B-Splines and does calculations in parallel with matrix operations. This computational method speeds up to 40% for large datasets. Moreover, Off-target effects can be added for general purpose usage.

We believe RNAiSpline assists and helps researchers to accelerate siRNA design.

## Abbreviations

AGO2: Argonaute-2
BiLSTM: Bidirectional Long Short-Term Memory
CNN: Convolutional Neural Network
GNN: Graph Neural Network
KAN: Kolmogorov-Arnold Networks
mRNA: Messenger RiboNucleic Acid
MLP: Multilayer Perceptron
MSE: Mean Squared Error
nM: Nanomolar
nt: Nucleotides
PCC: Pearson Correlation Coefficient
ReLU: Rectified Linear Unit
RNA: RiboNucleic Acid
RNA-FM: Ribonucleic Acid Foundation Model
RNAi: Ribonucleic Acid interference
IRSC: RNA-Induced Silencing Complex
ROC-AUC: Receiver Operating Characteristic – Area Under the Curve
RT-PCR: Reverse Transcription Polymerase Chain Reaction
siRNA: Small interfering Ribonucleic Acid

## Declarations

### Ethics approval and consent to participate

Not applicable

### Consent for publication Not applicable Availability of Data

The data underlying this article are available in the article.

### Code Availability and Implementation

The code for the RNAiSpline model is available at https://github.com/drugparadigm/RNAiSpline

### Author Contributions

**S.R.S**., **V.V.K**., **and S.S.S**.: Investigation; Conceptualization; validation; methodology; visualization; writing—original draft; formal analysis, data curation; **B.C.B**.: Methodology; validation; visualization; software; supervision; writing-review and editing. **V.K**.: Methodology; writing-review and editing; supervision.

## Acknowledgements

We authors, Sidhardha Reddy Surkanti, Vishnu Vardhan Kasturi, Sriram Shravan Saligram, Bhargava Chary Basangari, and Vani Kondaparthi, express our sincere gratitude to the Drugparadigm Research Lab for providing the necessary facilities and infrastructure that enabled the successful completion of this work.

